# Attention and control of posture: the effects of light touch on the center-of-pressure time series regularity and simple reaction time task

**DOI:** 10.1101/2024.04.12.589294

**Authors:** Anna Brachman, Justyna Michalska, Bogdan Bacik

**Affiliations:** Institute of Sport Sciences, Department of Biomechanics, The Jerzy Kukuczka Academy of Physical Education, 72a Mikolowska, Katowice, Poland; Institute of Sport Sciences, Department of Human Motor Behavior, The Jerzy Kukuczka Academy of Physical Education, 72a Mikolowska, Katowice, Poland

## Abstract

The stabilizing influence of a light touch on a postural sway has been consistently shown in the literature, however there is still no consensus in what way attentional resources are used when adopting additional tactile information during controlling an upright posture. To better elucidate the underlying mechanisms we introduced conditions of both sensory deprivation (closing the eyes), additional feedback (light touch), which seems to distracts from postural control and verified it by introducing dual task paradigm (i.e. measuring simple reaction time to an unpredictable auditory stimulus). Twenty five healthy students randomly performed eight postural tasks, four with (RT) and four without simple reaction task (NoRT). Center of pressure displacements were measured on a force plate in two visual conditions: eyes open (EO), eyes closed (EC) and two sensory conditions: with light touch (LT), without light touch (NoLT). Before each measurement participants were asked to consider the postural task as the primary task. Although simple reaction time did not differ between postural conditions (p>0.05), additional tactile information in anteroposterior direction caused decreased postural sway velocity (p<0.001, η2=0.86) and decreased standard deviation (p<0.001, η2=0.91) in both, reaction and visual conditions relative to NoLT conditions. Interestingly, simple reaction task modified subjects behavior in NoLT conditions and caused slower COP velocity (p<0.001, η2=0.53) without changes in signal regularity. Results also showed a significant increase in irregularity during standing with LT (p<0.001, η2=0.86) in both vision and reaction conditions, suggesting that the signal was more random. Although there were no significant changes between length of the reaction time between postural conditions but there was strong effect of light touch on COP regularity, we can conclude that light touch is attention demanding but changes of flow of attention are very subtle in this simple postural tasks. Furthermore COP regularity analysis is sensitive to even such subtle changes.

## 1. Introduction

During maintaining an upright posture, even in the absence of external perturbations, our body is in constant motion. It is well known that key sensory inputs for producing corrective torque attenuating postural sway via feedback control mechanisms come from visual, vestibular and proprioceptive sensory systems (1,2). The availability of sensory information impacts postural control. Removing visual information increase body sway whereas giving additional information provided by, for example, guiding eye movements to specific targets (3,4), providing haptic cues through light fingertip touch to a rigid surface or holding an anchor system leads to decrease in magnitude sway (5). The stabilizing influence of a light touch on a postural sway has been consistently shown in the literature (6–9). Haptic receptors supplies the postural control system with information about body’s relative position in space. It is an action-perception system which via exploratory behavior receives and integrates information about events that influence a subject (2,5). It has been suggested that this process is associated with increased use of attentional resources, presumably due to increased sensorimotor processing (10). Attention is defined here as the subjects’ limited capacity of the information processing. It is well known that performing any task requires some portion of this capacity (11). According to the resource competition hypothesis it is also said that in case of postural control and concurrent non-postural task, processes compete for a common processing resources. If an individual’s overall attention capacity is exceeded, performing an additional light fingertip touch task results in a processing capacity cost. This can lead to a degradation of one or both of given tasks (9,12,13).

There were few attempts to answer question whether light touch is attention demanding. Vuillerme et al. (8) have shown longer reaction responses to an unpredictable auditory stimuli during quiet standing performed with additional light touch when compared to baseline conditions while standing with eyes closed. Lee et al. (12) have shown that additional cognitive task (counting backwards) diminished stabilizing effect of light touch during standing. Authors concluded that these processes competed for finite pool of attentional resources. Also, it has been shown that cortical processing of sensory integration expressed by brain activity was increased in the posterior parietal cortex when standing with light touch (14). On the other hand there are several studies which have shown that additional cognitive task did not alter the stabilizing effect of the additional tactile information, thereby challenged the notion that this process requires attention. The stabilizing effect of the light touch was not deteriorated but actually it was stronger while performing a visual search task (searching for certain letters in a text) (9,15). Authors reported that postural sway can be adjusted to accomplish the other non-postural tasks, supporting the concept of functional integration and dismissing the theory of sensory competition. However, it’s worth noting that in these experiments, participants were not given any instructions on which task to prioritize. Additionally, they were not instructed to sway as little as possible, which suggests that the postural task might have been too simple to be affected by the concurrent non-postural task. The overall demands of the tasks may not have exceeded the participants’ available processing capacities. In another study by Barela et al. (16), it was observed that performing a cognitive task (backward counting) did not impact the stabilizing effect of light touch. The study concluded that enhanced sensory cues provided by light touch have little or no attentional demands. Therefore, there is still no consensus in what way attentional resources are used when adopting additional tactile information during controlling an upright posture.

The center of pressure (COP), as a variable, allows us to observe the actions of various neuromuscular components involved in maintaining upright posture. Its fluctuations provide valuable insights into the continuously active sensorimotor process of postural control (17–19). Over the past two decades, the use of nonlinear dynamics along with traditional balance measures has been an effective approach to gain additional insight into the mechanisms of postural control and further our understanding of the neuromuscular components involved (20).

To better elucidate the underlying mechanisms and attentional demands associated with taking advantage of additional tactile information, we combined traditional COP measures with regularity analysis and at the same time we introduced stimulus–response reaction-time protocol. Furthermore, participants’ focus of attention was controlled by giving them precise instructions. We hypothesized that light touch is attentional demanding hence it will cause increased reaction time length when compared to no light touch conditions. It will also bring increased level of randomness of the COP signal, suggesting more automatic control.

## 2. Materials and Methods

### 2.1. Participants

Twenty five healthy, active students (active no more than 6h per week) (17 women, 15 men; age: 22 ± 2 years (mean ± SD), body weight: 71 ± 14 kg, height: 175 ± 9 cm) from Academy of Physical Education in Katowice participated in this study. They were not aware about the purpose of the study. All participants gave written consent prior to participation required by the Declaration of Helsinki and the Bioethics Committee for Scientific Research at the Jerzy Kukuczka Academy of Physical Education in Katowice (no.1/2021). None of them had any current or previous neural, muscular or skeletal disorders that could influence the postural steadiness. Twenty three participants were right-handed and two left-handed with normal or corrected-to-normal vision. The limb dominance was stated by the participants and was consistent with the hand they used for writing.

### 2.2. Procedure

Participants performed four postural tasks and the same four postural tasks with simple reaction time task (dual task), overall subjects performed 8 postural conditions (Fig. 1). The order of the experimental conditions was randomized over subjects.

**Fig. 1.**
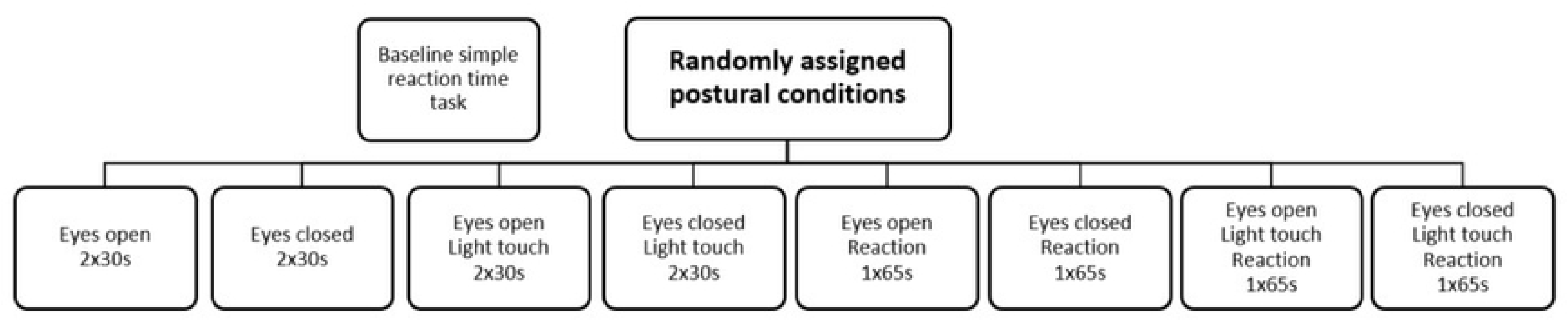
Experiment procedure.

Participants stood barefoot on a force plate (Kistler 9281C, USA), with feet placed parallel at hip width and big toes tangential to the line marked on the platform. The task was to sway as little as possible in two visual conditions: eyes open (EO), eyes closed (EC) and two tactile conditions: with light touch (LT), without light touch (NoLT). Each participant performed two 30s trials in each condition. During standing with additional tactile information (light touch) subjects stood with elbow of the dominant limb flexed approximately at 90° (8), wrist in the neutral position and the index fingertip of a hand in contact with the rigid surface which was flexibly attached to the tripod placed on the narrow support area. This solution prevented from getting a mechanical support for stance. Similar approach was adopted previously (9,21,22). Non-dominant arm hang naturally by the side. The position of the upper limbs in all other conditions was maintained and the tripod was removed from the subjects.

In order to verify the level of attention devoted to posture in each postural condition the dual task paradigm was introduced. Four postural conditions (EONoLT, ECNoLT, EOLT, ECLT) were conducted with additional simple reaction time (RT) task (Fig.1). During postural measurements subjects were asked to press their index finger, as quickly as possible, on a sensor placed on their thumb (non-dominant hand) to an unpredictable auditory stimulus. Prior to all measurements subject were instructed and trained how to react to a stimuli: “during all tasks keep your hand loose, after hearing the stimulus, press the sensor with your index finger and thumb as quickly as it is possible, then relax your hand again”. During the measurements the hand position and arrangement was controlled by the researcher. Subjects additionally performed a control trial in which their RT to five unpredictable auditory stimulus were measured in a relaxed standing position (10s trial). The control RT (baseline) was evaluated prior to all other conditions. No postural measurements were taken as this task only served to establish a baseline RT value for each subject.

In order to not to divert participants attention from postural task and to avoid influence of another motor task (pressing the sensor) on the COP fluctuations, each of the four conditions with additional reaction time task lasted 65s and auditory stimulus was randomly presented only once, between 30-33s of the trial. The data between 30s and 35s were extracted from future analysis to avoid the influence of startle effect on the COP fluctuation. Furthermore, the data before (first 30s of the trial) and after stimuli (second 30s of the trial) were compared to verify whether the RT influenced the COP fluctuations (23) (Fig.2). Similarly like in previous research (8) before each measurement participants were asked to consider the postural task as the primary task, thus changes in RT across conditions presumably would reflect changes in the attentional resources necessary for performing the postural task.

**Fig. 2.**
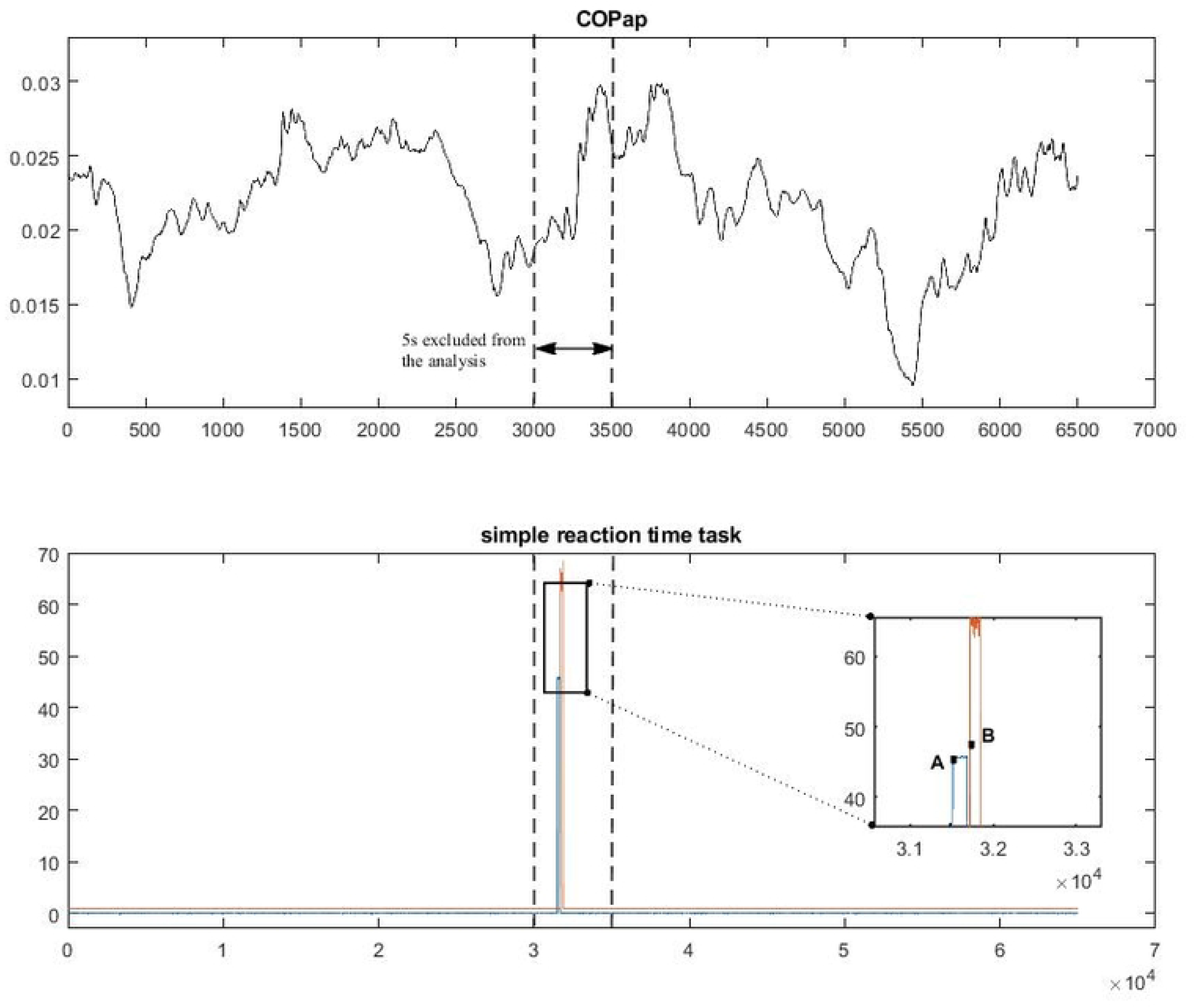
Typical COP fluctuations in anteroposterior direction during quiet standing with RT task (upper panel) and reaction time which was determined by temporal interval between the onset of the auditory stimulus (A) and the subjects’ response (B) (lower panel).

### 2.3. Apparatus and data analysis

We decided to analyze influence of the light touch in sagittal plane as in this plane control must be more constrained due to more degrees of freedom of the kinematic chain. All postural measurements were taken on the force platform (Kistler, MA, USA) with sampling frequency 1000Hz. During the RT task, a special set was used that was verified in previous study (24). It consisted of a tactile sensor (Patent PL 222119.B1) mounted on the thumb, connected to a signal generator module (Patent PL 222753.B1) coupled to the sensor of the DTS EMG system (Noraxon, Scottsdale, AZ, USA). The acoustic stimulus (sound f=1000Hz, lasting about 200ms) was also recorded by transmitting directly the analog signal from the sound generator to the A/D input of the measurement card (Kistler 5691A, sampling frequency 1000Hz). All signals (acoustic stimulus, subject’ s response and the force plate data) were synchronized.

The reaction time (in milliseconds) was defined as the temporal interval between the onset of the auditory stimulus and the subjects’ response (pressing the sensor) (Fig.2).

For the amplitude COP analysis signal was filtered with bidirectional, 2^nd^ order low pass Butterworth filter with 12.5Hz cut off frequency and down sampled by a factor of 10 to obtain 100Hz frequency. Two variables were examined, mean velocity (cm/s), and standard deviation (cm) indexing variability or the amount of postural sway.

For the COP regularity analysis signal was down sampled but was not filtered as filtering could remove the inherent properties of the signal (25,26). Regularity of the signal was evaluated by means of Sample entropy (27). The signal was also inspected according to its stationarity and detrended, if needed. To optimize the length of a template vector (m) and tolerance level (r) the minimization of the maximum entropy relative error method was implemented (27) In the current study m = 2 and r = 0.05 x SD were adopted. SampEn is the negative natural logarithm for conditional probability that a series of data points within a certain length, m, would be repeated within the length m + 1. The higher level of Sample entropy express higher randomness of the time series (27).

All procedures and signal processing methods were performed offline using MATLAB r2017b software (Mathworks Inc., Natick, MA, USA).

### 2.4. Statistical analysis

To analyze influence of light touch on a simple reaction time a one-way repeated –measures analyses of variance (ANOVA) was applied. To verify influence of dual task (simple reaction task) on COP fluctuations before and after presentation of the auditory stimuli the dependent t-test or Wilcoxon test (according to normal distribution assumption) was applied in each evaluated variable and each vision and tactile condition. As there were no differences before and after stimuli presentation, mean values of two 30s trials from RT conditions were used for further analysis. A three-way repeated-measures ANOVA (2×2×2) with two levels of vision (EO vs. EC), two levels of tactile condition (NoLT vs. LT) and two levels of reaction task (NoRT vs. RT) was used for statistical comparison of different conditions. When the Mauchly’s sphericity assumption was violated, the more conservative Geisser-Greenhouse F-test was performed. Level of significance was set at 0.05. Post-hoc analyses (Newman-Keuls) were used when a significant main effect was observed. All statistical analyses were conducted using Statistica software package (TIBCO Software Inc., USA) version 13.1.

## 3. Results

### 3.1 Reaction time

Results showed significant influence of postural conditions on the reaction time task (F(4,96)= 13.03, p<0.001 η2=0.31). Reaction times were significantly longer when compared to baseline, relaxed standing condition (p<0.001) (Fig. 3). However, there were no additional significant differences between all other conditions.

**Fig. 3.**
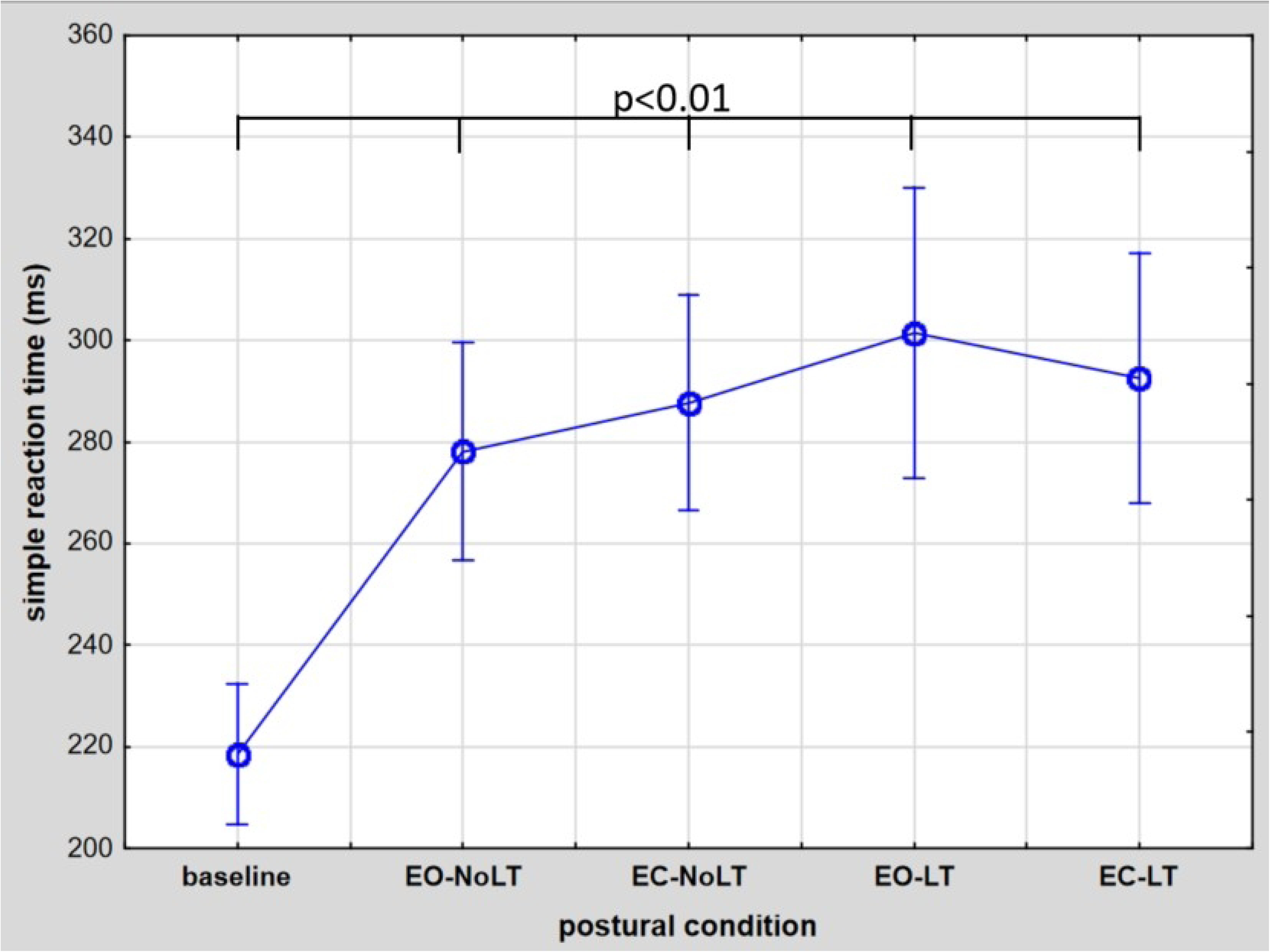
Mean reaction time obtained for the different postural conditions. Baseline – relaxed standing position, EO – eyes open, EC – eyes closed, NoLT – quiet standing without light touch, LT - quiet standing with light touch. Error bars represents 95% confidence intervals.

### 3.2 Influence of RT on *before* and *after* COP fluctuations

Statistical analysis showed no significant influence of reaction time task on the COP fluctuations before and after auditory stimuli in all COP parameters (p>0.05) (Fig. 4,5,6).

**Fig. 4.**
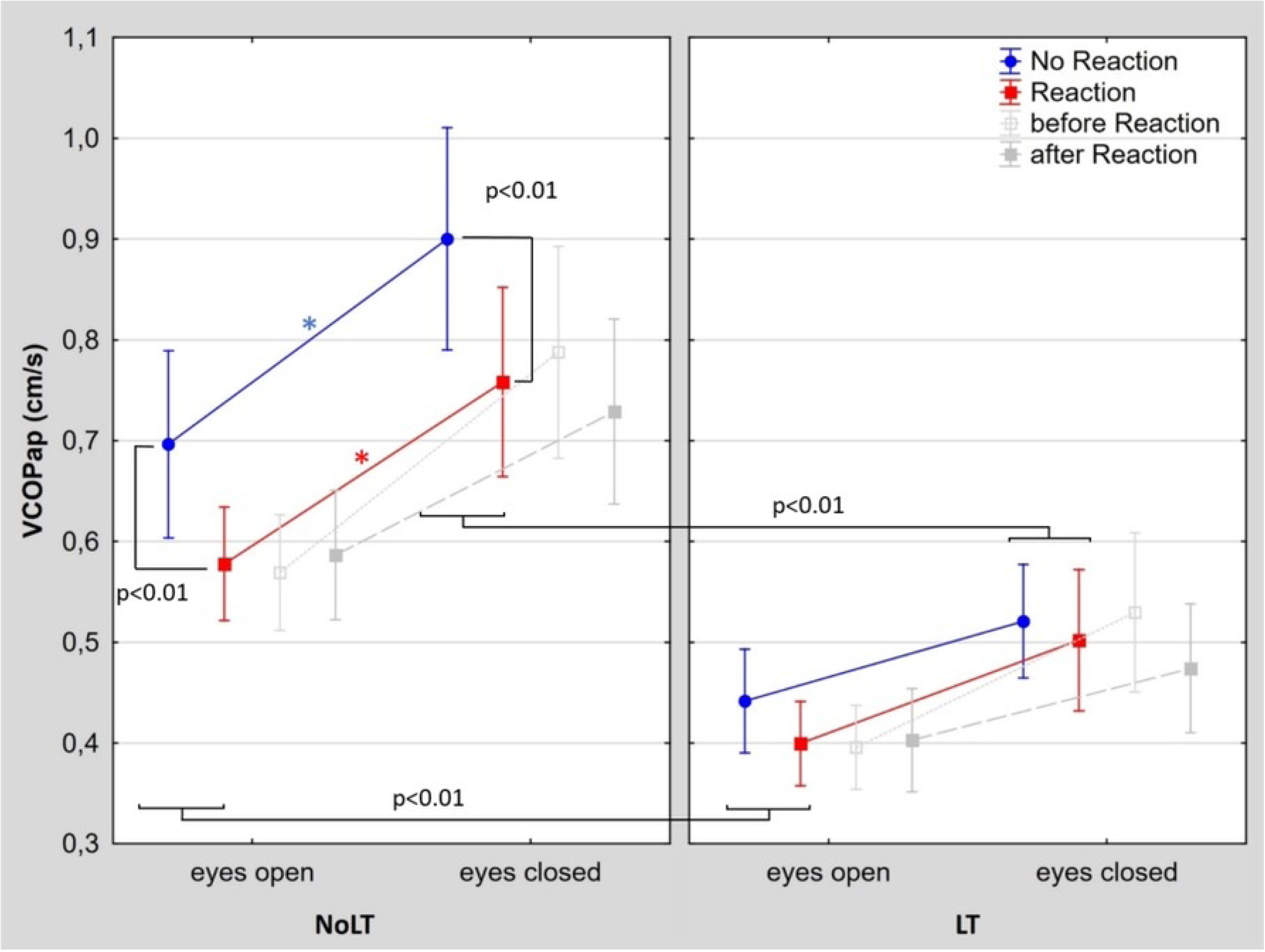
Mean COP Velocity observed during different postural conditions. EO – eyes open, EC – eyes closed, NoLT – quiet standing without light touch, LT - quiet standing with light touch. Error bars represent 95% confidence intervals. * indicate significant differences between vision conditions (p < 0.05). Grey squared marks show results *before* and *after* auditory signal presentation during reaction task conditions. Red squared marks represent mean values of *before* and *after* signal presentation taken for further analysis.

**Fig. 5.**
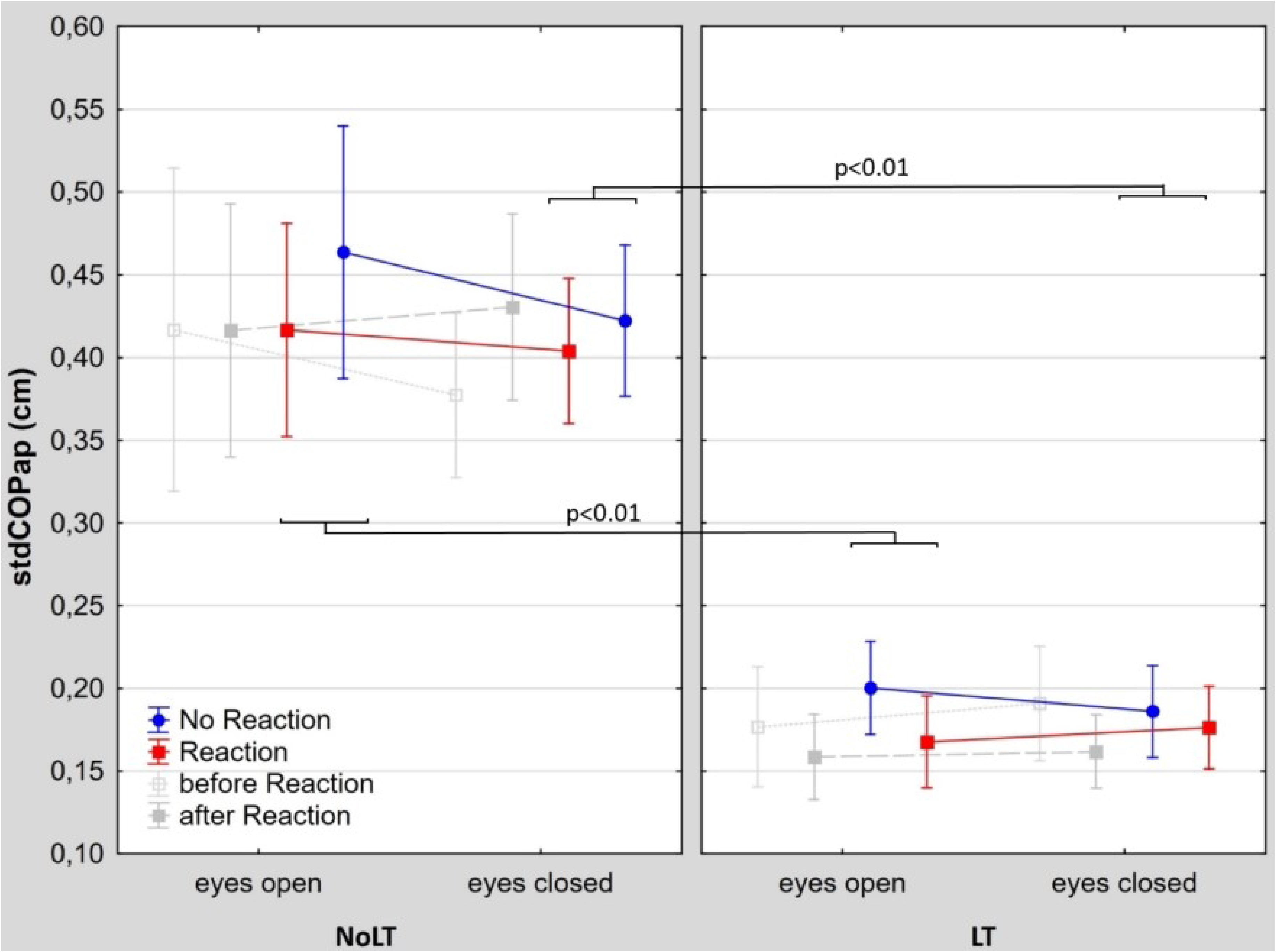
Mean COP standard deviation observed during different postural conditions. EO – eyes open, EC – eyes closed, NoLT – quiet standing without light touch, LT - quiet standing with light touch. Error bars represent 95% confidence intervals. Grey squared marks show results *before* and *after* auditory signal presentation during reaction task conditions. Red squared marks represent mean values of *before* and *after* signal presentation taken for further analysis.

**Fig. 6.**
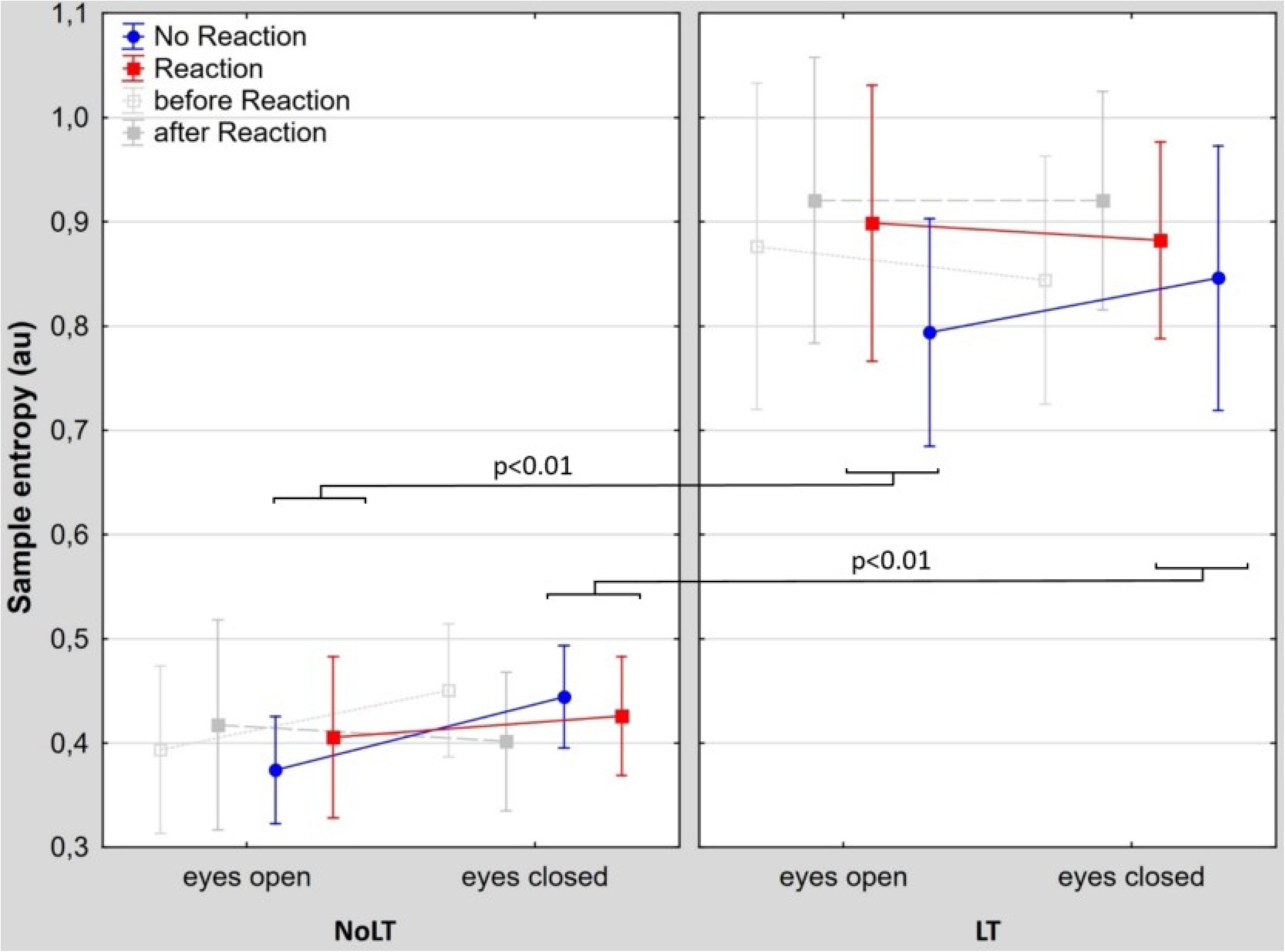
Mean COP regularity level (sample entropy) observed during different postural conditions. EO – eyes open, EC – eyes closed, NoLT – quiet standing without light touch, LT - quiet standing with light touch. Error bars represent 95% confidence intervals. Grey squared marks show results *before* and *after* auditory signal presentation during reaction task conditions. Red squared marks represent mean values of *before* and *after* signal presentation taken for further analysis.

### 3.3 COP velocity

Vision and tactile condition significantly affected sway velocity in sagittal plane (F(1,24)=35.8, p<0.001 η2=0.59, F(1,24)=149.4, p<0.001 η2=0.86, respectively); sway velocity was significantly higher in EC when compared to EO condition in both tactile and reaction task conditions (Fig. 4). Light touch caused significant decrease of sway velocity in both vision conditions. However, significant vision × tactile condition interaction (F(1,24)=9.41, p<0.01 η2=0.28) showed that light touch dampened the effect of vision deprivation. The reaction task also significantly affected sway velocity (F(1,24)=27.15, p<0.001 η2=0.53). The post hoc analyses showed that the velocity was lower when subjects had to react for the auditory stimuli, in both vision conditions (p<0.001). The tactile condition × reaction task interaction effect was also significant (F(1,24)=6.41, p=0.018 η2=0.21) indicating that light touch decreased sway velocity to such extent that velocity did not differ between RT and NoRT (EO p= 0.06, EC p=0.38).

### 3.4 Standard deviation

Tactile condition significantly affected the variability of the COP position in the anteroposterior direction (F(1,24)=270.7, p<0.001 η2=0.91) (Fig. 5). Light touch caused significant reduction of the sway variability in both vision conditions. There was also significant reaction task effect (F(1,24)=4.7, p=0.04 η2=0.16), however post-hoc analysis did not reveal significant differences, only in EONoLT difference was close to significant when compared RT to noRT conditions (p=0.057). No other main and interaction effects were significant.

### 3.5 Sample entropy

Sample entropy in the anteroposterior direction was significantly affected by the tactile condition (F(1,24)=153.3, p<0.001 η2=0.86) (Fig. 6). The additional tactile information increased COP irregularity in both vision and reaction task conditions. No other main and interaction effects were significant.

## 4. Discussion

The main goal of the present study was to examine whether attentional demands associated with the use of additional tactile information is increased during an upright standing in various postural conditions in healthy young adults. Furthermore, our aim was to verify whether light touch influences predictability of the COP fluctuations in these conditions. In agreement with many previous studies (6–9), present results revealed that light touch had stabilizing effect on postural control, it caused decreased postural sway velocity and decreased sway variability in the sagittal plane, in both vision and reaction conditions. Surprisingly, contrary to our hypothesis, participants did not significantly lengthen their simple reaction time for an unpredictable auditory stimuli, which at first glance would seem that LT was not attention demanding. These results are in contrast to that obtained by Vuillerme et al. (8). Previous authors (for review see: Woollacott et al. (11)) highlighted that one of the main challenges for the reliability of the dual task paradigm is shifting attention or sacrificing quality of the main task to improve outcomes in the secondary task (here, postural stability and reaction task respectively). The first threat in the current study was that participants would change their postural behavior before and after given unpredictable auditory stimuli, but it was not the case. The constancy of the postural task quality (before and after reaction) was confirmed statistically. However, the results revealed that despite the precise instruction given before each trial (to sway as little as possible and to always prioritize the postural task), subjects’ behavior in NoLT conditions was influenced by reaction task. While waiting for the sound participants were even more stable than without RT task, they were swaying slower. Possible explanation to these results is that performing a RT task created a so called *alert state* (23,28). Even though participants during NoRT postural tasks were consciously trying to be as stable as possible, during reaction conditions they were prepared for an action production (pressing the button), hence unconsciously decreased postural sway velocity even more. This interpretation is corroborated by previous studies of giving invalid cues produced prior to a postural perturbation which created *generalized alert state* and resulted in a decreased muscle onset latencies (29). In current study changes in subjects’ postural control were present not only while waiting for the perturbation (it would resemble anticipatory adjustment mechanism), this *alert state* was still present after “perturbation”, so participants somehow adopted different strategy to control posture in reaction conditions. Previously, Vuillerme et al. (23) examined influence of reaction time task on postural control but their results were partially different. Authors reported decreased postural sway velocity, but only during reaction production (i.e. participants verbally indicated colour of the presented light) and after reaction production when compared to control group and values before the reaction production. In current study we observed decreased postural sway velocity both, before and after reaction production, when compared to NoRT conditions. Discrepancies of these results can stem from methodological differences. Vuillerme et al. (23) compared 3 very short COP temporal data sets (5s) from 25s trials. Furthermore, the stimulus given to the subjects (visual) and reaction production (verbal) were different than in the current study. However, the question, why in the current study participants changed behavior before and after stimuli and in Vuillerme et al. (23) study participants did not reveal changes while waiting for the stimuli when compared to control group, remains unknown.

In the current study the presence of light touch diminished the influence of reaction conditions, possibly because of a floor effect, with additional tactile information participants could not be more stable (i.e. sway slower, decrease variability). Previous authors postulated that performing a RT task lead to increased level of attention and/or muscle stiffness. The question is whether in current study participants shifted their attention towards secondary task (reaction production) or increased stiffness. The question is not trivial. According to results obtained by previous authors (15,30,31) increasing stiffness should increase velocity of the postural sway, thus this mechanism does not explain behavior of our subjects. We cannot directly answer this question. However, based on the explanation of the COP regularity proposed by previous authors (32–35), the lack of significant changes in the sample entropy in this conditions (RT vs. NoRT) would suggest that changes in sway velocity were not due to the participants’ attention being drawn towards the reaction task. Moreover, according to the resource competition hypothesis (11), devoting more attention to a reaction task than a postural task should impair the latter. In this case, no such deterioration but further improvement was observed. It is possible that participants decreased the exploration of the environment and purposeful migration of the reference position of the body (18,30,36). This does not necessarily mean that at the same time feedback control, which brings more regular COP signal and has a higher processing cost, was decreased. This indicates that postural fluctuations are related to constraints imposed by the tasks and can be controlled to facilitate performance of non-postural tasks (here reaction task) without changing the level of automaticity (21,37,38). We also assume that the processing cost (amount of attention) related to additional tactile information is small and/or the probe-reaction task is insufficient to such subtle changes.

We expected that sample entropy might shed some light on the mechanisms responsible for the changes caused by additional tactile information. Indeed, in the sagittal plane results showed a significant increase in irregularity during standing with LT in both vision and reaction conditions, suggesting that the signal was more random and participants relied more on a feedforward control (i.e. signal stem from more automatic control). Thus, the level of attention devoted to control posture decreased. Our findings are supported by a prior research on the effect of LT on the COP regularity level (13). The study indicated that LT was responsible for increase in sample entropy. What is interesting, these results are similar even though authors adopted different data processing before sample entropy calculations and did not adjusted input parameters (vector length, tolerance level) to their data. Taking into account the fact that sample entropy is sensitive to the choice of input parameters and signal processing (26,39,40) it seems that sample entropy is very sensitive to even small differences in COP fluctuations.

It is known that dual task has impact on postural control (11,30). Current results suggests that waiting for the auditory stimuli influenced subjects postural control, this is limitation of the current study. Another limitation is lack of the lower leg muscle activity observation, which would help to answer question whether participants increased stiffness during reaction conditions and would help to better elucidate causes of the observed COP fluctuation changes. Another studies with more challenging postural conditions and additional analysis, like COP signal decomposition (41) or sway density plots (18) could further increase insights into changes in postural control mechanisms related to simple reaction time task and additional haptic information.

## 5. Conclusions

Although there were no significant changes between length of the reaction time between postural conditions generally we can conclude that light touch is attention demanding but changes of flow of attention are very subtle in this simple postural tasks. Furthermore COP regularity analysis is sensitive to even such subtle changes.

## Acknowledgments

We would like to acknowledge Sobota Grzegorz for customizing the tactile sensors and reaction task set [24] to the specific purpose of this study.

## References

1. Horak FB, Henry SM, Shumway-Cook A. Postural perturbations: new insights for treatment of balance disorders. Phys Ther. 1997;77(5):517–33.

2. Peterka RJ, Goodworth AD, Mellodge P, Peterka RJ, Volpe D, Giantin MG, et al. Sensorimotor Integration in Human Postural Control. J Neurophysiol. 2002;1097–118.

3. Giveans MR, Yoshida K, Bardy B, Riley M, Stoffregen TA. Postural sway and the amplitude of horizontal eye movements. Ecol Psychol. 2011;23(4):247–66.

4. Figueiredo GA, Barela AMF, Bonnet CT, Barela JA. Haptic Information Improvement on Postural Sway is Information-Dependent But Not Influenced by Cognitive Task. J Mot Behav. 2022;54(4):515–22. 10.1080/00222895.2022.2030667

5. Mauerberg-Decastro E, Moraes R, Tavares CP, Figueiredo GA, Pacheco SCM, Costa TDA. Haptic anchoring and human postural control. Psychol Neurosci. 2014;7(3):301– 18.

6. Jeka JJ, Lackner JR. Fingertip contact influences human postural control. Exp Brain Res. 1994;79(2):495–502.

7. Riley MA, Stoffregen TA, Grocki MJ, Turvey MT. Postural stabilization for the control of touching. Hum Mov Sci. 1999;18(6):795–817.

8. Vuillerme N, Isableu B, Nougier V. Attentional demands associated with the use of a light fingertip touch for postural control during quiet standing. 2006;232–6.

9. Chen FC, Chen HL, Tu JH, Tsai CL. Effects of light touch on postural sway and visual search accuracy: A test of functional integration and resource competition hypotheses. Gait Posture. 2015;42(3):280–4. 10.1016/j.gaitpost.2015.06.001

10. Ishigaki T, Ueta K, Imai R, Morioka S. EEG frequency analysis of cortical brain activities induced by effect of light touch. Exp Brain Res. 2016;234(6):1429–40.

11. Woollacott M, Shumway-Cook A. Attention and the control of posture and gait: A review of an emerging area of research. Gait Posture. 2002;16(1):1–14.

12. Lee Y, Goyal N, Aruin AS. Effect of a cognitive task and light finger touch on standing balance in healthy adults. Exp Brain Res. 2018;236(2):399–407. 10.1007/s00221-017-5135-9

13. Lara JR, da Silva CR, de Lima FF, da Silva MC, Kohn AF, Elias LA, et al. Effects of light finger touch on the regularity of center-of-pressure fluctuations during quiet bipedal and single-leg postural tasks. Gait Posture. 2022;96:203–9.

14. Ishigaki T, Imai R, Morioka S. Cathodal transcranial direct current stimulation of the posterior parietal cortex reduces steady-state postural stability during the effect of light touch. Neuroreport. 2016;27(14):1050–5.

15. dos Santos DG, Prado-Rico JM, Alouche SR, Garbus RB de SC, de Freitas PB, de Freitas SMSF. Combined effects of the light touch and cognitive task affect the components of postural sway. Neurosci Lett. 2019;703:99–103.

16. Barela AMF, Caporicci S, de Freitas PB, Jeka JJ, Barela JA. Light touch compensates peripheral somatosensory degradation in postural control of older adults. Hum Mov Sci. 2018;60:122–30. 10.1016/j.humov.2018.06.001

17. Collins SL, De Luca. A language for software subsystem composition. Exp Brain Res. 1993;308–18.

18. Baratto L, Morasso PG, Re C, Spada G. A new look at posturographic analysis. Motor Control. 2002;6:246–70.

19. Maurer C, Peterka RJ. A new interpretation of spontaneous sway measures based on a simple model of human postural control. J Neurohysiology. 2005;93:189–200.

20. Riley MA, Turvey MT. Variability and Determinism iln Motor Behavior. J Mot Behav. 2002;34(2):99–125.

21. Riley MA, Wong S, Mitra S, Turvey MT. Common effects of touch and vision on postural parameters. Exp Brain Res. 1997;117(1):165–70.

22. Brachman A, Sawicka A, Bednarz B, Akbas A, Słomka K. The effect of a light finger touch on the signal complexity during quiet standing. Acta Bioeng Biomech. 2022;24(4).

23. Vuillerme N, Nougier V, Teasdale N. Effects of a reaction time task on postural control in humans. Neurosci Lett. 2000;291:77–80.

24. Żak M, Mikrut G, Sobota G. Laboratory and Natural Environment Condition. Sensors (Basel). 2023;23:1–11.

25. Rhea CK, Silver TA, Hong SL, Ryu JH, Studenka BE, Hughes CML, et al. Noise and complexity in human postural control: Interpreting the different estimations of entropy. PLoS One. 2011;6(3):1–9.

26. Lubetzky A V., Harel D, Lubetzky E. On the effects of signal processing on sample entropy for postural control. PLoS One. 2018;13(3):1–15.

27. Lake DE, Richman JS, Pamela Griffin M, Randall Moorman J. Sample entropy analysis of neonatal heart rate variability. Am J Physiol - Regul Integr Comp Physiol. 2002;283(3) 52–3.

28. Kuczyński M, Szymańska M, Bieć E. Dual-task effect on postural control in high-level competitive dancers. J Sports Sci. 2011;29:539–45.

29. McChesney JW, Sveistrup H, Woollacott MH. Influence of auditory precuing on automatic postural responses. Exp Brain Res. 1996;108(2):315–20.

30. Yamagata M, Falaki A, Latash ML. Effects of voluntary agonist–antagonist coactivation on stability of vertical posture. Motor Control. 2019;23(3):304–26.

31. Warnica MJ, Weaver TB, Prentice SD, Laing AC. Gait & Posture The influence of ankle muscle activation on postural sway during quiet stance. Gait Posture. 2014;39(4):1115–21. 10.1016/j.gaitpost.2014.01.019

32. Roerdink M, De Haart M, Daffertshofer A, Donker SF, Geurts ACH, Beek PJ. Dynamical structure of center-of-pressure trajectories in patients recovering from stroke. Exp Brain Res. 2006;174(2):256–69.

33. Roerdink M, Hlavackova P, Vuillerme N. Center-of-pressure regularity as a marker for attentional investment in postural control: A comparison between sitting and standing postures. Hum Mov Sci. 2011;30(2):203–12. 10.1016/j.humov.2010.04.005

34. Stins JF, Michielsen ME, Roerdink M, Beek PJ. Sway regularity reflects attentional involvement in postural control: Effects of expertise, vision and cognition. Gait Posture. 2009;30(1):106–9.

35. Krȩcisz K, Kuczyński M. Attentional demands associated with augmented visual feedback during quiet standing. PeerJ. 2018;(6):1–11.

36. Latash ML, Ferreira SS, Wieczorek SA, Duarte M. Movement sway : changes in postural sway during voluntary shifts of the center of pressure. 2003;314–24.

37. Fraizer E V, Mitra S. Methodological and interpretive issues in posture-cognition dual-tasking in upright stance. Gait Posture. 2008;27(2):271–9.

38. Chen FC, Tsai CL. The mechanisms of the effect of light finger touch on postural control. Neurosci Lett. 2015;605:69–73. 10.1016/j.neulet.2015.08.016

39. Yentes JM, Hunt N, Schmid KK, Kaipust JP, McGrath D, Stergiou N. The appropriate use of approximate entropy and sample entropy with short data sets. Ann Biomed Eng. 2013;41(2):349–65.

40. Brachman A, Sobota G, Bacik B. The influence of walking speed and effects of signal processing methods on the level of human gait regularity during treadmill walking. BMC Sports Sci Med Rehabil. 2022;14(1):1–11. 10.1186/s13102-022-00600-4

41. Zatsiorsky VM, Duarte M. Instant Equilibrium Point and its Migration in Standing Tasks Rambling and.pdf. Motor Control. 1999;3:28–38.

